# Seven amino acids gate transcriptional activation by a minimal MarA Helix-turn-helix DNA-binding domain

**DOI:** 10.64898/2026.09.23.753698

**Authors:** Marina Corbella, Ariadna Serrano, Roberto Bello-Madruga, Marc Torrent, Ivan Erill, Jessica M.A. Blair, Enea Sancho-Vaello

## Abstract

Prokaryotic transcription factors (TFs) lie at the core of antimicrobial resistance, controlling genes that let bacteria survive antibiotic exposure. The AraC/XylS family of TFs is defined by a ∼99-residue DNA-binding domain composed of two helix-turn-helix (HTH) motifs. While this two-motif architecture is considered the minimal functional unit, the striking sequence and structural similarity between both HTH motifs raises the question of whether it evolved from a single ancestral HTH domain. Here, we designed two C-terminal truncations of MarA, comprising a single HTH motif, differing by seven-residues (IRSRKMT). Electrophoretic-mobility shift assays reveal that both constructs specifically bind the marbox sequence as reconstituted dimers, while size-exclusion chromatography shows they exist in a monomer-dimer equilibrium in solution. Despite retaining DNA-binding capacity, the truncations diverge functionally: while MarA64 (including IRSRKMT) activates transcription and confers regular erythromycin tolerance, MarA57 (lacking IRSRKMT) yields a transcriptionally inactive complex that suppresses reporter expression below baseline, suggesting competitive promoter occupancy. Molecular dynamics simulations and AlphaFold models suggest that the IRSRKMT extension forms an α-helical element stabilizing a transcriptionally productive dimer interface. Conversely, its loss disrupts quaternary assembly, alters DNA bending, and misaligns RNA polymerase-contacting residues. Furthermore, free dimers explore non-productive conformations, suggesting that functional dimerization occurs upon DNA engagement. These findings establish that a single, correctly dimerized HTH domain is sufficient for both DNA binding and transcriptional activation, providing a structural rationale for short AraC/XylS-like proteins and offering a tuneable scaffold for synthetic biology and novel anti-virulence strategies.

## Introduction

Transcription factors (TFs) constitute a central layer of control in the cell, governing which genes are expressed, when, and to what extent. In bacteria, this control is exerted largely at the level of transcription initiation, through proteins that bind specific or semi-specific DNA sequences and either promote or block recruitment of RNA polymerase to a promoter (Browning and Busby 2016). Through this mechanism, TFs modulate the selective synthesis of proteins essential to processes ranging from nutrient metabolism to stress survival, biofilm formation, or cell attachment, and, in pathogenic organisms, the expression of virulence factors (Roncarati and Scarlato 2017; Henderson et al. 2021; Mancera et al. 2021). Because a single TF can control multiple, functionally-related genes, disrupting its DNA-binding or activation function, can collapse an entire physiological program rather than a single gene product, making TFs attractive both as tools for engineering gene expression and as targets for antimicrobial strategies aimed at virulence or resistance pathways rather than viability itself (Roncarati et al. 2022). Understanding the molecular mechanisms underlying selective DNA binding and productive transcriptional activation is therefore a prerequisite for designing new or modified transcription factors capable of deliberately tuning the cell’s defensive or adaptive capacity.

Among transcriptional regulator families, the AraC/XylS family is an interesting target for structural and mechanistic studies for several reasons. First, it is large and near-ubiquitous in Bacteria: members were identified in 80.1% of prokaryotic genomes, and together with the LysR and TetR/ArcR families it accounts for roughly 30% of all bacterial TFs identified to date, while remaining entirely absent from Eukaryotes (Cortes-Avalos et al. 2021). Second, the family harbours members regulating genes involved in general metabolism, stress response, and virulence, spanning organisms ranging from environmental bacteria to major human pathogens (Gallegos et al. 1997; Cortes-Avalos et al. 2021). Third, this functional breadth is matched by substantial architectural diversity, despite all members sharing a single conserved DNA-binding fold (Gallegos et al. 1997; Cortes-Avalos et al. 2021).

The family-defining feature is a ∼99-residue DNA-Binding Domain (DBD) with two helix-turn-helix (HTH) motifs separated by an α-helix, folding into seven conserved α-helices (H1–H7; with H1–H3 defining HTH1, H5–H7 defining HTH2, and H4 acting as a linker; Figure 1A). Despite sharing this common domain, the family exhibits a wide architectural diversity. In a recent review on the family, 65.4% of analysed proteins were multidomain, carrying 1 to 15 additional companion domains (CD) (Cortes-Avalos et al. 2021). These CDs typically serve sensory functions (e.g., arabinose for AraC, aromatic compounds for XylS) and/or mediate dimerization, with their position varying between N-terminal, C-terminal, and internal placements across different members. The remaining members (34.6%) are monodomain, consisting only of the DBD (*E. coli* SoxS and MarA are standard examples of this minimal architecture (Rhee et al. 1998; Shi et al. 2022b)). Notably, the same study identified 73 proteins shorter than the ∼90 residues required to accommodate the full seven-helix DBD, but still accommodating both HTH motifs, which the authors flagged as requiring experimental follow-up rather than treating as confirmed functional variants (Cortes-Avalos et al. 2021). Whether such truncated architectures can support DNA binding, let alone transcriptional activation, has not yet been tested.

**Figure 1:**
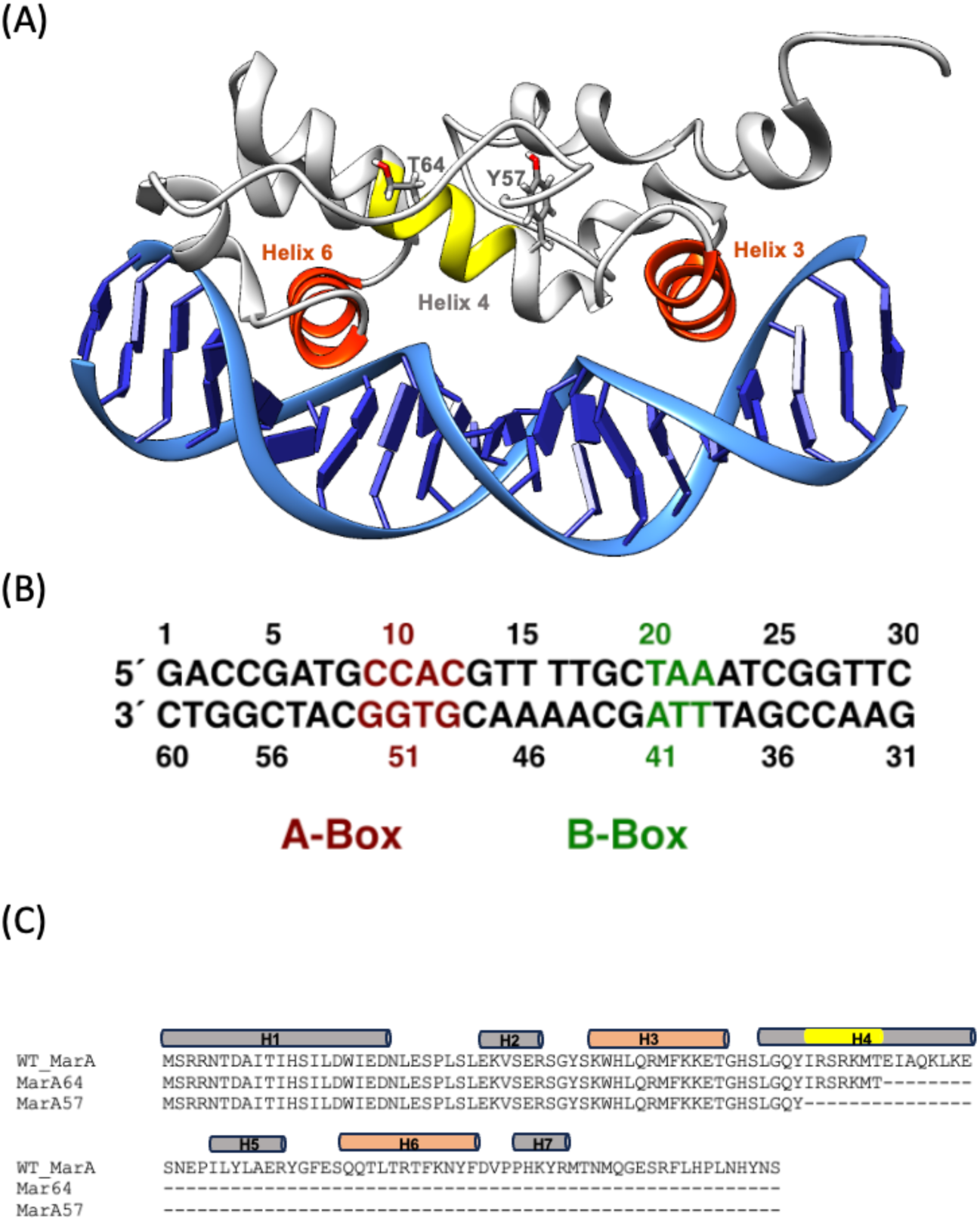
**(A)** Structure of the WT MarA–DNA complex (Rhee et al. 1998), with truncation boundaries for MarA64 and MarA57 indicated by residues T64 and Y57. The 7-residue fragment present in MarA64 but absent in MarA57 is shown in yellow. Helix 3 and Helix 6 (orange), each part of a helix-turn-helix motif, make direct contact with the DNA. **(B)** The marbox sequence corresponding to *marRAB* operon is shown with A- and B- boxes coloured in maroon and green, respectively. **(C)** Sequence alignment of WT MarA, MarA64, and MarA57, showing the IRSRKMT extension present in MarA64 but absent in MarA57. Secondary structure elements (H1–H7, coloured as in (A)), annotated above the WT MarA sequence, are based on PDB 1XS9 (Dangi et al. 2004).

Global regulators, such as MarA, SoxS, and Rob, occupy a distinct tier within bacterial transcriptional networks (Martinez-Antonio and Collado-Vides 2003; Duval and Lister 2013). Unlike regulators controlling a single operon, global regulators bind numerous promoters (often dozens to over a hundred) coordinating the expression of genes that may belong to functionally unrelated pathways (metabolism, stress response, virulence, efflux) in response to an environmental or physiological stimulus. This broad reach is frequently paired with a relaxed, degenerate recognition sequence, since a single strict consensus site would be incompatible with binding so many distinct promoter contexts (Martinez-Antonio and Collado-Vides 2003).

MarA (multiple antibiotic resistance protein A), the global multidrug-resistance regulator of *Escherichia coli*, is an AraC/XylS monomeric TF consisting of two HTH domains connected by a linker helix, with no separate regulatory module (Figure 1A). By using two helices (H3 and H6), it binds a single degenerate 20-bp DNA sequence called the marbox at over 40 promoters in *E. coli*, including the *acrAB* operon encoding the principal AcrAB-TolC multidrug efflux pump (Okusu et al. 1996; Martin and Rosner 2011). The marbox is composed of two half-sites: the A-box, contacted by helix 3 of the N-terminal HTH domain, and the B-box, contacted by helix 6 of the C-terminal HTH domain (Rhee et al. 1998) (Figure 1A). Structural, biochemical, and computational studies all point to the A-box as the primary driver of DNA binding, while sequence variations at the B-box exert a comparatively minor impact on binding affinity (Li and Demple 1996; Kwon et al. 2000; Corbella et al. 2021).

The MarA N-terminal helix has been also shown to be involved in Lon degradation, which terminates activation of downstream genes, and in RNA polymerase binding (Griffith et al. 2004). In addition, we have recently shown that the dynamics of this helix are directly involved in the protein’s ability to bind DNA (Corbella et al. 2025). Since MarA has no ligand-binding domain and activates transcription exclusively through direct RNAP contact (Dangi et al. 2004), any MarA truncation retaining DNA binding but losing activation must have disrupted either the RNAP interface or the geometry required to present it correctly.

Structural alignment of the two MarA HTH domains reveals substantial conservation at the DNA-contacting positions (Arg44/Arg94, Lys47/Lys97, and Ser38/Ser88 are structurally equivalent across the two domains) suggesting that they may have arisen by gene duplication of a common ancestral single-HTH unit. This raises the possibility that an isolated HTH unit, presented as a dimer, could retain sufficient information to bind the marbox and induce transcription on its own.

To address this question, we designed two C-terminal truncations of the MarA N-terminal HTH domain differing by only seven residues. Using a combination of *in vitro* binding assays, *in vivo* reporter systems, and enhanced sampling molecular dynamics (MD) simulations, we evaluated their stoichiometry, DNA-binding capacity, and transcriptional output. We show that these two constructs exhibit opposing functional profiles despite retaining sequence-specific marbox occupancy, revealing how subtle C-terminal modifications alter dimerization geometry and RNA polymerase recruitment. Ultimately, these findings demonstrate that a single, correctly dimerized HTH domain is sufficient to drive both DNA binding and transcriptional activation, redefining the minimal functional architecture of the AraC/XylS transcription factor family.

## Results and Discussion

### Two N-terminal MarA truncations specifically bind the marbox

To define the minimal structural unit required for marbox recognition and to evaluate whether DNA binding is sufficient for transcriptional activation, we designed a C-terminal truncation of MarA based on the high sequence conservation at the DNA-contacting positions across both HTH domains: Arg44/Arg94, Lys47/Lys97, and Ser38/Ser88 (numbering according to Uniprot code P0ACH5). Truncation MarA(1-64) (hereafter referred to as MarA64) was designed to encompass half of the wild-type (WT) MarA protein, cutting helix 4 at position Thr64 (Figure 1).

After overexpression of WT MarA and MarA64 in *E. coli* T7 Express cells and purification by Ni-NTA affinity chromatography (Figure S1), their ability to bind DNA was tested by electrophoretic mobility shift assays (EMSA). Both, WT MarA and MarA64 were able to bind the *marRAB* and *acrAB* marboxes using 200-bp (Figure 2A) or 30-bp (Figure 2B) DNA fragments including the sequence of these marboxes in the center of the fragment (Table S1), although MarA64 showed lower DNA affinity and required higher protein concentrations (600– 800 ng) to visualise the binding. None of the two proteins were able to shift a 30-bp non-specific DNA fragment, showing that MarA64 maintains the sequence specificity for DNA binding (Figure 2B). Unexpectedly, the DNA-bound complex formed by MarA64 migrated to a position comparable to that of WT MarA. Given that the monomeric mass of MarA64 is roughly half that of WT MarA, this raised the hypothesis that MarA64 assembles into a dimer on DNA.

**Figure 2.**
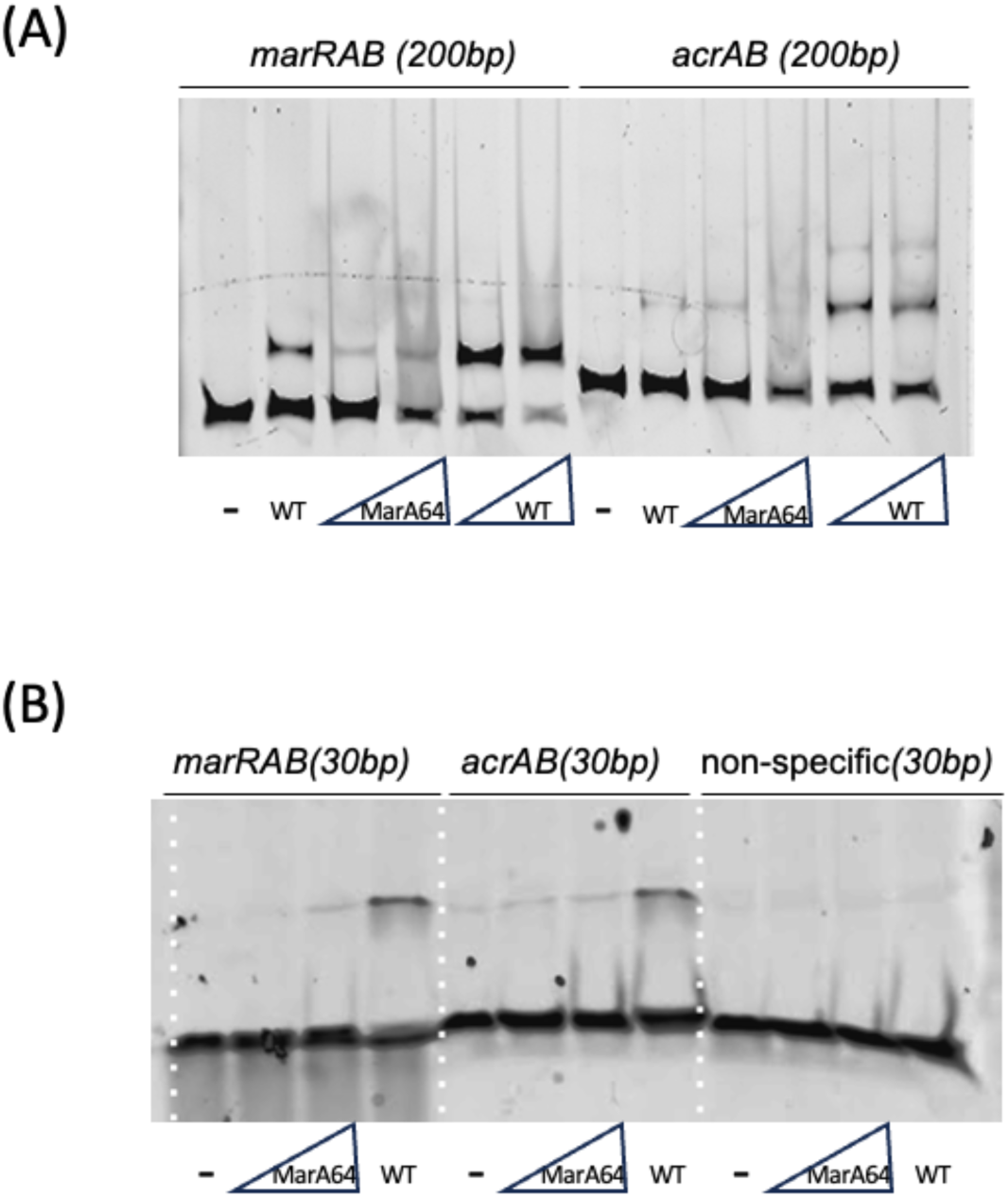
EMSA of WT MarA and MarA64 binding to **(A)** 200-bp or **(B)** 30-bp DNA fragments containing the *marRAB* or *acrAB* marboxes. A 30-bp non-specific DNA fragment was included as a specificity control. For each construct, protein was titrated as indicated by the triangle (increasing concentration) or added at a single fixed concentration (single lane), with a no-protein control (–) in each set. In both, **(A**) and **(B)**, WT MarA was tested at 1, 2 and 2.5 µM, and MarA64 at 5 and 6 µM. The weaker shifted-band intensity for MarA64 relative to WT MarA, suggests reduced DNA-binding affinity.

Intrigued by the observed apparent dimer formation, we used AlphaFold3 (Abramson et al. 2024) to model MarA64 dimer-DNA complex. The overall structure was below the confidence threshold (ipTM = 0.34, pTM = 0.48), but the prediction was assembled as a dimer with a MarA-like conformation (Figure 3A). Based on the predicted dimer interface, we designed a new construct, MarA(1-57) (hereafter referred as MarA57), truncated at Tyr57 to remove the C-terminal IRSRKMT tail (ipTM = 0.28pTM = 0.36) (Figure 1C and 3A). The rationale behind this seven-residue deletion was to shorten the C-terminus, thereby repositioning helix 1 of the second monomer to mimic helix 4 of wild-type MarA and ultimately favouring the assembly and stabilization of the dimeric DNA-bound complex.

**Figure 3.**
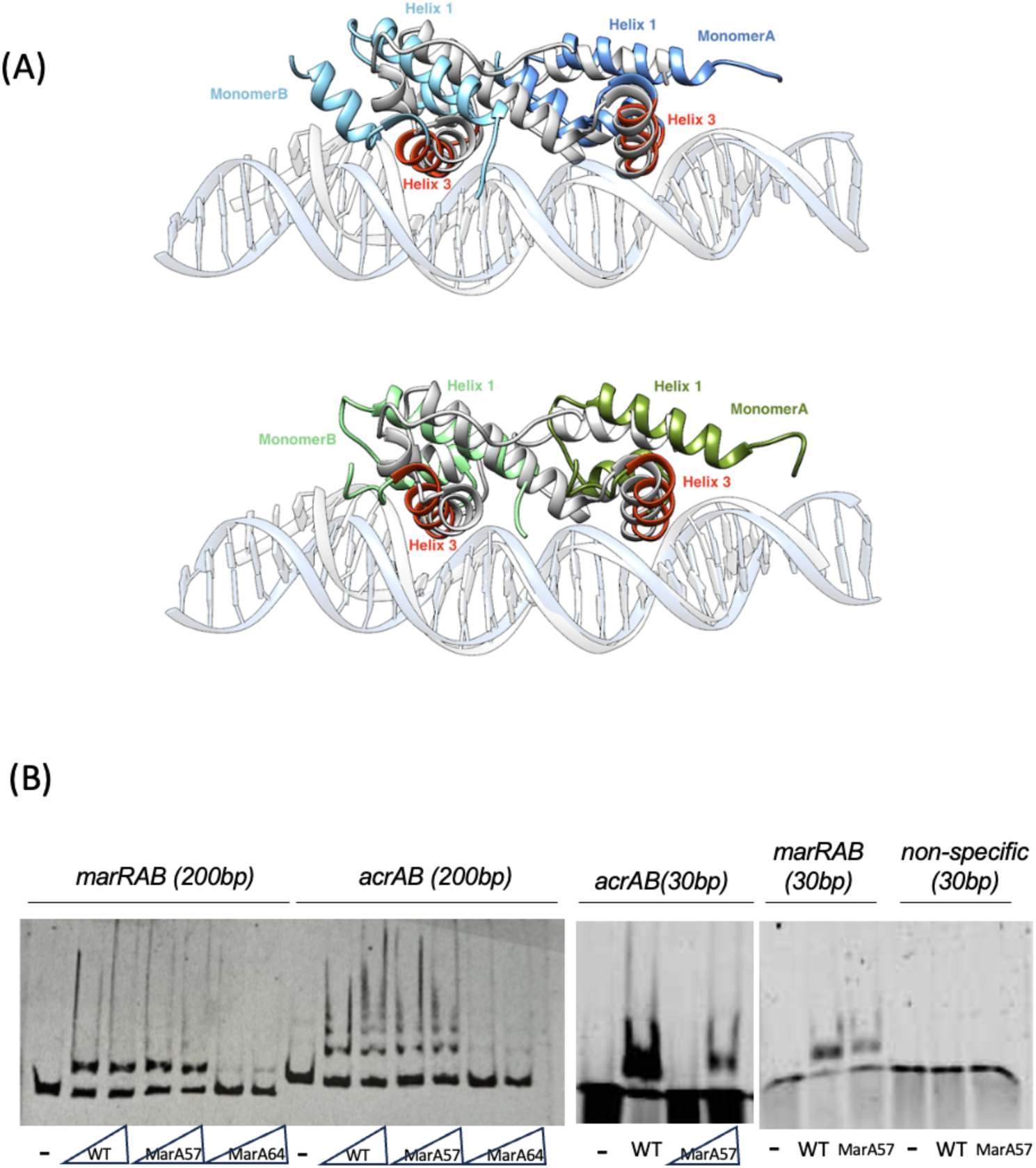
(A) Overlay of WT MarA (gray, PDB ID: 1BL0 (Rhee et al. 1998)) with AlphaFold3 models for MarA64 (blue, top) and MarA57 (green, bottom). Each monomer is depicted with a different shade of either blue or green, while helixes 3 are shown in orange. Monomers are named A or B related to the marbox they bind. **(B)** EMSA of WT MarA, MarA57, and MarA64 binding to 200-bp fragments containing the *marRAB* or *acrAB* marbox (left), or 30-bp fragments containing the *acrAB* marbox, the *marRAB* marbox, or a non-specific sequence (right). For each construct, protein was titrated as indicated by the triangle (increasing concentration) or added at a single fixed concentration (single lane), with a no-protein control (–) in each set. WT MarA was tested at 1, 2 and 2.5 µM, and Mar57 and MarA64 at 5 and 6 µM. The weaker shifted-band intensity for MarA64 relative to MarA57, suggests reduced DNA-binding affinity.

EMSA assays with the MarA57 truncation showed a band shift at a position similar to that of MarA64 and WT MarA. MarA57 displayed higher affinity than MarA64 for both the *marRAB* and *acrAB* marboxes on the 200-bp fragments, albeit still lower than WT MarA (Figure 3B). As with MarA64, MarA57 retained sequence-specific DNA binding, showing no substantial shift of the non-specific 30-bp fragment (Figure 3B). Notably, MarA64 produced a less complete shift than MarA57, suggesting that removal of the IRSRKMT extension increases, rather than decreases, marbox affinity *in vitro*.

### MarA truncations exist in a monomer–dimer equilibrium in solution

The electrophoretic mobility of both truncation-DNA complexes in EMSA, was similar to that of the WT MarA–DNA complex, despite the truncations being approximately half the molecular weight (∼9.8 kDa or ∼8.9 kDa versus ∼17.4 kDa). This is consistent with a 2:1 stoichiometry (two truncation monomers per DNA fragment), which reconstitutes a complex of comparable mass to the 1:1 WT MarA-DNA complex.

To determine complex stoichiometry, we performed isothermal titration calorimetry (ITC). However, the protein concentrations required to yield interpretable thermograms exceeded the solubility/stability limits of WT and truncated MarA, in buffers compatible with their native folds, preventing the acquisition of reliable binding isotherms.

To clear out the oligomerisation state of free WT MarA and both truncations, we performed HPLC with a size-exclusion column (Figure 4). WT MarA eluted as a single, sharp peak at ∼9.6 min. Both truncations displayed this same early-eluting peak, but additionally showed a second, broader, tailing peak at ∼11.7 min that was absent in WT MarA. As larger species elute earlier than smaller ones in size-exclusion chromatography, we interpret the early, shared peak as the higher-order (dimeric truncations, as well as monomeric WT MarA) species and the later, truncation-specific peak as the corresponding monomer, consistent with a monomer/dimer equilibrium for both truncated constructs that is not populated, or not resolved, for WT MarA under these conditions. This assignment is consistent with the 2:1 (dimer:DNA) stoichiometry inferred from EMSA (Figures 2 and 3). In addition, the presence of these species is supported by the slightly different time of elution for the three peaks, being the first to elute the MarA64 dimer (19512 Da), followed by the MarA57 dimer (17767 Da) and lastly WT MarA monomer (17347 Da) (Figure 4). The broad, tailing shape of the monomer peak, in contrast to the sharp early cluster, are consistent with secondary (hydrophobic) interactions with the stationary phase retarding the monomer beyond ideal size-exclusion behaviour. Notably, the monomer peak was markedly larger in relative area for MarA57 than for MarA64, indicating a shift in population toward the monomeric state for MarA57.

**Figure 4.**
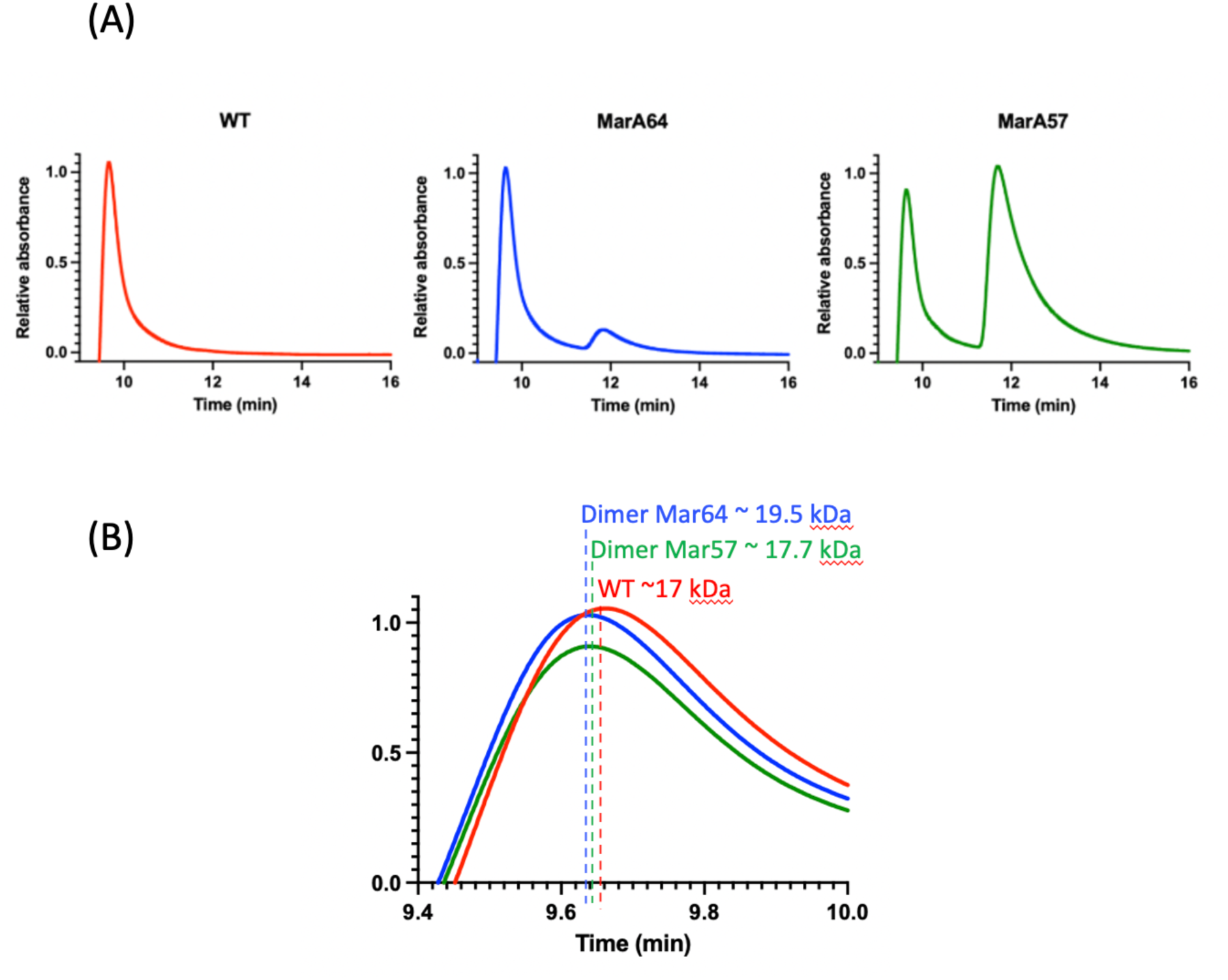
**(A)** Size-exclusion chromatograms (Phenomenex bioZen 1.8 µm SEC-2, 150 × 4.6 mm; ammonium acetate pH 6.8; detection at 220 nm) of WT MarA (red), MarA64 (blue), and MarA57 (green). Buffer was subtracted and data were normalised to 1 for each run. **(B)** Zoomed in on the first peak to show the different elution times of the three proteins. They eluted in the order expected from the sequence-predicted masses of the truncation dimers and the WT MarA monomer.

### A seven amino acids truncation switches transcriptional output

To evaluate whether these truncations could drive transcriptional activation in addition to DNA binding, both of them were assayed using a reporter system in which the *acrAB* promoter, including the marbox, drives *gfp* expression in *E. coli* T7 Express cells (Corbella et al. 2025). Interestingly, the two truncations exhibited opposite functional profiles. Upon IPTG induction, MarA64 increased GFP fluorescence above the empty vector baseline, albeit less efficiently than WT MarA. In contrast, MarA57 consistently yielded GFP fluorescence below the empty vector baseline under IPTG-induced conditions, a suppression phenotype not observed for either MarA64 or WT MarA (Figure 5).

**Figure 5.**
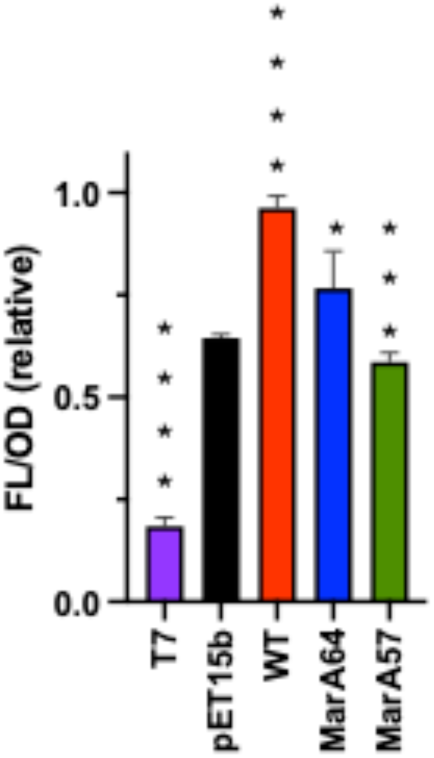
Reporter assay: normalized GFP fluorescence (fluorescence/OD600) for pET15b (black), or pET15b harbouring the gene encoding WT MarA (red), MarA64 (blue), or MarA57 (green). T7 cells (purple) normalised GFP fluorescence is also included. Samples were IPTG-induced. The asterisks indicate statistically significant differences between pET-15b empty and the rest of the strains (tested by a Student’s t test).

These results suggest that MarA64 can functionally mimic the MarA-RNAP contact region (Dangi et al. 2004) when bound to the DNA, consistent with the structural model generated by AlphaFold (Figure 3A). Conversely, MarA57 presents an interesting picture. It binds DNA *in vitro* but suppresses transcription *in vivo.* This suppression phenotype may suggest competitive marbox occupancy by a transcriptionally inactive species: MarA57 occupies the *acrAB* marbox, excludes endogenous MarA, and presents an altered RNAP interaction surface that either cannot make productive contact or is geometrically misaligned, despite initial AlphaFold predictions (Figure 3A).

### Seven amino acid C-Terminal truncation disrupts the dimerization interface

Building on the observation that removal of the IRSRKMT sequence in MarA57 alters transcriptional output despite its high structural similarity to the WT MarA-DNA complex (PDB ID: 1BL0 (Rhee et al. 1998); Figure 3A), we sought to get dynamical molecular insights into the structural changes induced by both C-terminal truncations. To address this, we used the AlphaFold-generated 2:1 models to perform molecular dynamics simulations on both complexes (3 x 2.5 μs). Note that although the overall structures were below the confidence threshold, the protein fold, and specifically the contacting helices H3, display confident pLDDT > 70 (Figure S2).

A central question is why transcriptional activation is affected in MarA57 while DNA binding is preserved. The most parsimonious interpretation would be that the altered oligomerization geometry of these variant places key RNAP-contacting residues in a configuration incompatible with productive RNAP recruitment, while still allowing marbox occupancy. Consistent with this hypothesis, analysis of the MD trajectories revealed tight insertion of both HTH motifs within the corresponding marbox (Figure 6A), alongside distinct conformational behaviour within the dimer interface in the C-terminal truncated complex (Figure 6B,C).

**Figure 6.**
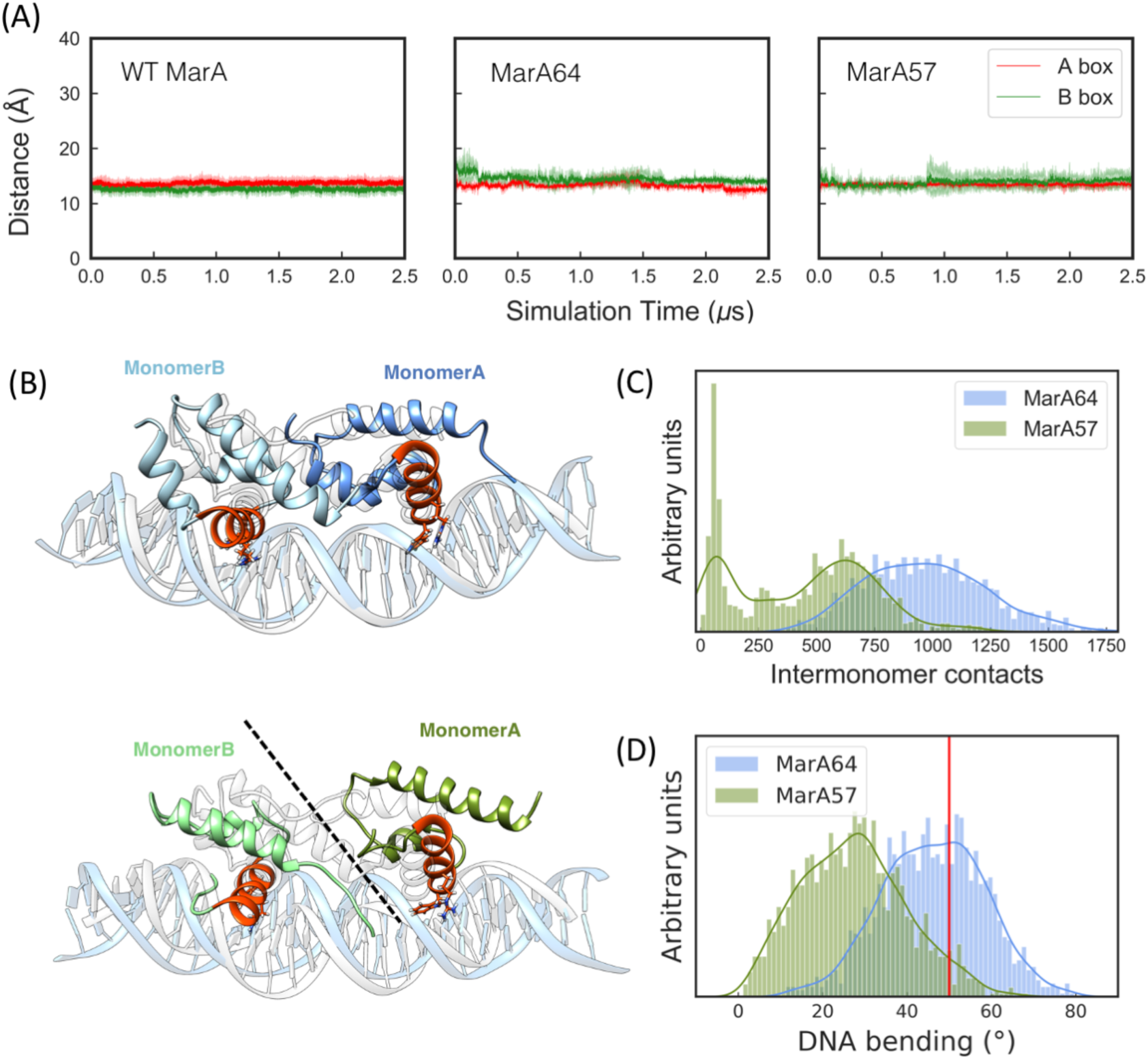
**(A)** Time evolution of the distances between helix 3 inserted in the major groove and the base pairs at the A- and B-boxes (shown in red and green, respectively). **(B)** Structures of the most populated clusters of MarA64-*mar* (top) and MarA57-*mar* (bottom). WT MarA x-ray structure PDB ID 1BL0 (Rhee et al. 1998) is overlayed with MarA64-*mar* (top) for structural comparison. **(C)** Distribution of the number of contacts between monomers A and B. D) Distribution of the DNA bending angle (details in Materials and Methods). The red vertical line indicates the average DNA bending angle along WT MarA MD simulation. All the data was extracted along 3 x 2.5 μs of MD simulation of WT MarA-*mar* (Corbella et al. 2021), MarA64*- mar and* MarA57*-mar* complexes.

As expected, both truncations retain the capacity to bind the marbox by inserting their respective helix 3 elements into the A- and B-boxes. Both truncations comprise two units of the N-terminal DNA-binding domain of MarA, which harbour key residues driving tight DNA binding (Arg44 and Trp40). The crucial impact on DNA binding of these two residues has been previously reported, both computationally and experimentally, via mutations introduced either into the residues themselves or into the interacting nucleobases (Gillette et al. 2000; Rousseau et al. 2004; Taliaferro et al. 2012; Corbella et al. 2021). This stability is evident when tracking the distances between residues Lys39-Thr50 of helix 3 and their target base pairs in the A- and B-boxes (nucleotide ID 10-11, 48-49 for A box and 19-20, 41-42 for B box) (Corbella et al. 2021) (Figure S3), with both truncations displaying MarA comparable distances (Figure 6A). Alongside these primary interactions, Trp40 establishes persistent T-shaped π-p interactions with C9/10 in the A box, and C19 in the B box (Table S2), as well as persistent hydrogen bonding between Arg44 and G8/G51/G52 for the A box, and G18/G42 for the B box (Table S3).

Remarkably, removal of the C-terminal IRSRKMT sequence disrupts the dimerization interface between the two monomers, drastically compromising the number of intermonomer key stabilizing contacts (Figure 6B,C). In MarA64, this C-terminal IRSRKMT sequence forms an α-helical element (residues Leu54 to Thr64) that enables critical stabilizing interactions with helix 1 of the adjacent monomer (Figure S4). Consequently, the lack of this structured region in MarA57 destabilizes the inter-subunit interface and leads to the loss of the characteristic MarA-like quaternary assembly. Crucially, this compromised dimeric architecture directly impacts the conformation of the bound DNA (Figure 6D). In the functional WT MarA-DNA complex, the induced specific DNA bending angle is strictly required to properly orient the protein for RNAP recruitment (Rhee et al. 1998; Dangi et al. 2004; Shi et al. 2022a; Shi et al. 2022b). However, the relative orientation of the HTH domains in MarA57 fails to induce this native curvature, imposing instead an aberrant DNA bending angle (Figure 6B, D). This distortion could ultimately misalign the critical RNAP-contacting surface necessary for transcriptional initiation.

### Free dimers explore non-productive configurations

To investigate the conformational dynamics of the free truncated variants and determine whether dimeric assembly occurs before or after DNA binding, we performed Protein-Protein Interaction-Gaussian Accelerated Molecular Dynamics (PPI-GaMD) simulations on both MarA64 and MarA57. Although initial AlphaFold predictions of the free dimers did not suggest a MarA-like preorganization (Figure S5), we initiated our simulations from MarA-like dimeric assemblies derived from dimeric MarA64- and MarA57-DNA complexes to test the stability of a preorganized dimer. Interestingly, both systems rapidly dissociate from these preorganized starting states and rebound after a few nanoseconds, exploring a highly heterogeneous conformational space. However, only a single dissociation/rebinding event was captured across each of the six independent replicates, which precluded robust kinetic estimations of dimer assembly (Wang and Miao 2022).

Detailed examination of the PPI-GaMD trajectories revealed that the free dimers adopted alternative quaternary arrangements structurally incompatible with marbox recognition and binding (Figure 7). The absence of unique canonical dimeric structures complicated the selection of specific interface descriptors. Therefore, we used generic descriptors like the center-of-mass (COM) distance between monomers and the total number of inter-subunit contacts to map the conformational landscape of both systems. Clustering the combined trajectories of each system using RMSD allowed us to identify preferred dimeric assemblies for both variants and project them on the 2D landscape (Figure 7). Notably, the most populated cluster for MarA64 accounted for 51% of the simulation time, whereas the primary cluster for MarA57 represented only 24% (Figure S6). This trend aligns well with our SEC experiments, which showed that the monomer–dimer equilibrium is shifted towards dimer formation for MarA64 (Figure 4).

**Figure 7.**
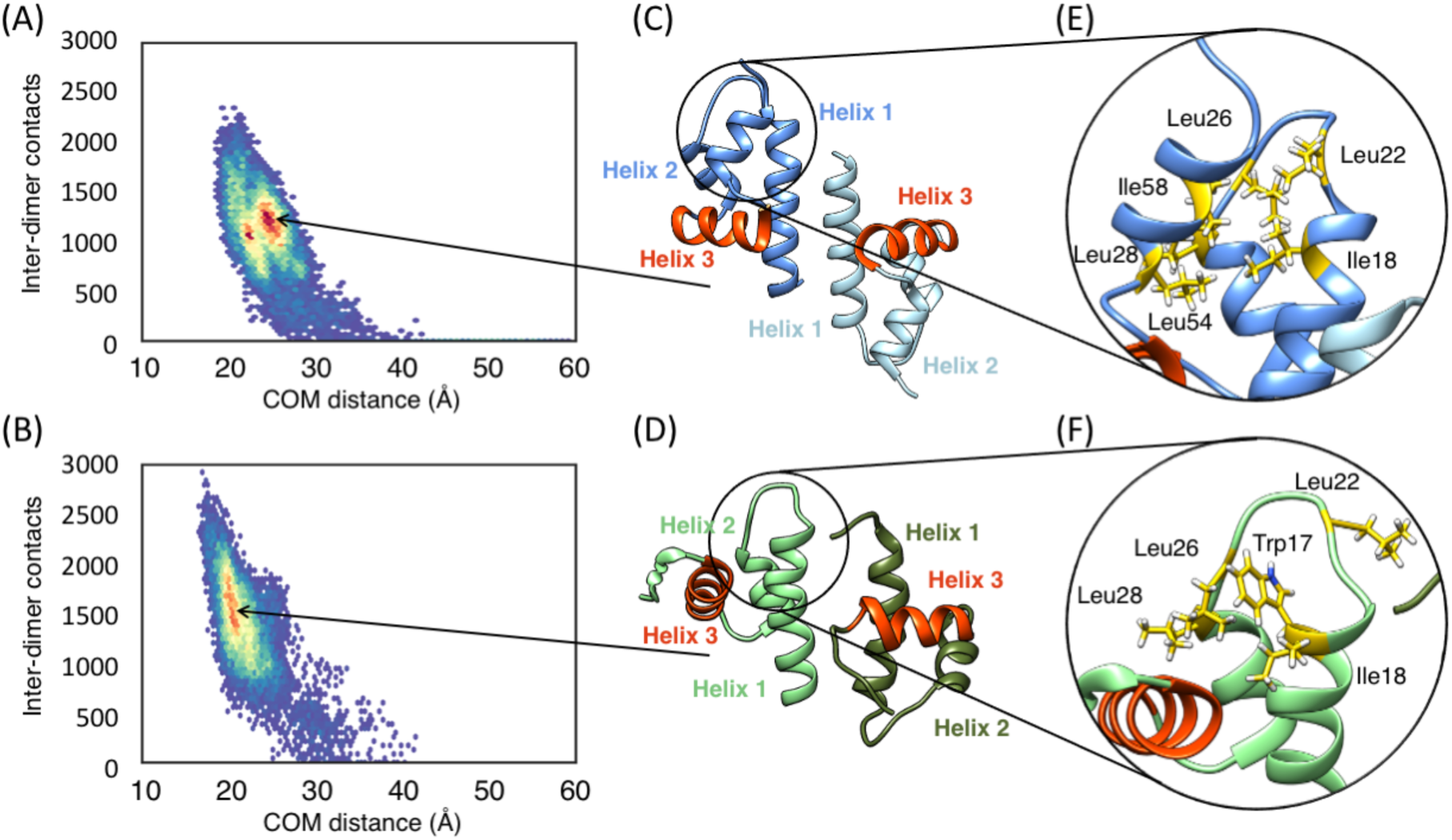
2D histogram of the COM distance between monomers vs. the number of inter-dimer contacts in free **(A)** MarA64 and **(B)** MarA57 dimers, alongside with the structure of the most populated clusters **(C)** and **(D)**, respectively. Close-up of the hydrophobic loop between helix 1 and helix 2 of free **(E)** MarA64 and **(F)** MarA57 dimers. The hydrophobic residues from the loop (Trp17, Ile18, Leu22, Leu26 and Leu28) are shown in yellow sticks, as well as the hydrophobic residues (Leu54 and Ile58) from the α-helical C-terminus that pack against it in MarA64 dimer. Each monomer is colored in a different shade of blue (MarA64) or green (MarA57), while helices 3 are coloured in orange. Data was extracted along 6 independent 2 μs PPI-GaMD simulations.

Both systems tend to engage through helix 1-helix 1 contacts. In MarA64, the C-terminal IRSRKMT sequence forms an α-helical element that packs against a hydrophobic loop between helix 1 and helix 2, as helix 4 in WT-MarA (Figure 7 and S6, S7), thereby stabilizing this transcriptionally non-productive dimeric conformation. Contrarily, the C-terminal truncation in MarA57 exposes the hydrophobic patch, rendering the system more prone to either bind DNA to shield this region or form transient dimeric species.

These computational findings directly reconcile our experimental SEC and EMSA observations. SEC shows that MarA57 shifts the monomer–dimer equilibrium predominantly toward the monomeric species, whereas MarA64 favours dimer formation. Crucially, EMSA assays revealed superior DNA-binding affinity for MarA57 variant. Combined with the structural instability observed in both PPI-GaMD and AlphaFold models, these results suggest a model in which the monomer is the primary DNA-binding competent species, with functional dimerization occurring only upon marbox engagement. In this framework, pre-formed dimers in solution may represent inactive states incapable of productive marbox engagement, which would explain why shifting the equilibrium toward monomers enhances DNA binding.

### Expression of MarA57 is associated with reduced growth under erythromycin stress

The AcrAB-TolC efflux pump is the principal tripartite multidrug efflux system in *Enterobacteriaceae* and is transcriptionally activated by MarA through direct induction of both the *acrAB* operon and *tolC*. To first assess whether MarA truncations altered antibiotic susceptibility, we determined the minimal inhibitory concentration (MIC) for erythromycin and ciprofloxacin, two well-characterised AcrAB-TolC substrates in overnight cultures of strains overexpressing WT MarA, MarA64, or MarA57, and found no clear differences among the three strains for either antibiotic (512 ug/ml for pET empty vector; > 512 µg/ml for the other strains). Because MIC determination by serial dilution has limited resolution — differences are only detectable in discrete two-fold steps — we reasoned this assay might not capture subtler, efflux-dependent effects on antibiotic tolerance. To test whether overexpression of WT MarA, MarA64, or MarA57, was instead associated with altered growth under antibiotic stress consistent with AcrAB-TolC-dependent tolerance, we monitored growth of *E. coli* T7 Express cultures carrying pET15b empty vector (pET), WT MarA, MarA64, or MarA57 at two different erythromycin concentrations. To allow adaptation of the cultures, they were grown to OD_600_=0.5 and induced with 0.5 mM IPTG for 3 h at 37°C prior to the growth-curve assay. To sustain MarA overexpression and plasmid selection during the growth curve, 0.5 mM IPTG and 50 µg/mL ampicillin were maintained.

Even when the strains were not exposed to erythromycin, cultures overexpressing WT MarA or MarA64 showed a shorter apparent lag phase relative to the empty-vector control, with MarA57 showing an intermediate growth onset. Because this baseline difference in growth onset was independent of antibiotic exposure, we decided to compare strains at a fixed timepoint (8 h) rather than at full saturation, to capture differences during active growth before any construct reached plateau. At this timepoint, MarA57 reached a lower OD600 than WT MarA, and MarA64, at every erythromycin concentration tested (Figure S8). This pattern was not found when using gentamicin as antibiotic, a non-AcrAB-TolC efflux substrate (data non shown). This finding is consistent with WT MarA and MarA64 retaining, and MarA57 losing, a MarA-dependent function associated with erythromycin tolerance.

### Gene duplication as an evolutionary strategy to improve DNA affinity

The structural conservation between the N- and C-terminal HTH domains of MarA (Figure S9A), combined with our results suggesting that two isolated HTH domains can dimerize to bind DNA and induce transcription, led us to hypothesize that a gene duplication event of an ancestral single-HTH domain may have increased DNA affinity, with helix 4 emerging as a key element in correctly positioning the two independent HTH domains in the orientation and geometry required for correct DNA binding and transcriptional induction. If this hypothesis holds, remnants of isolated MarA-like HTH domains might still be present in extant genomes.

To test this, we built a Hidden Markov Model (HMM) profile of the MarA HTH1 using a multiple sequence alignment of MarA, Rob, and SoxS, and high-coverage homologs identified via BLASTP outside the *Enterobacteriaceae* (Figure S9B). We then used the MarA HTH1 profile (Supplemental Data 1) to search the UniProt Bacteria (taxID: 2) dataset. This search yielded 700 protein sequences shorter than full-length MarA (127 aa), and 87 sequences shorter than the MarA DBD (99 aa) (Supplemental Data 2). We examined the AlphaFold-predicted structures for a subset of these sequences, mapping to complete protein records with lengths (38-60 aa) comparable to the MarA HTH1 motif (43 aa), and we found them to match the predicted structure of our isolated HTH domains (Figure S9). In this way, the computational search identified candidate proteins consistent with isolated MarA-like HTH domains; whether these proteins are functional or capable of binding DNA remains to be tested experimentally.

### Conclusions

In this study, we demonstrate that a single N-terminal HTH domain of MarA is sufficient to reconstitute sequence-specific DNA binding and drive transcriptional activation. Our findings indicate that the minimal functional architecture of the AraC/XylS family can be significantly shorter than the canonical two-motif topology (Cortes-Avalos et al. 2021). Crucially, we identify a seven-residue C-terminal sequence (IRSRKMT) that acts as a structural switch dictating the functional outcome of the truncated transcription factor. While its presence in MarA64 stabilizes the dimeric geometry required for productive RNAP recruitment and subsequent transcriptional activation, its removal in MarA57 yields a transcriptionally inactive complex that could competitively occupy the promoter.

Furthermore, our biophysical and computational analyses provide a plausible mechanistic view of the assembly pathway for these regulatory complexes. PPI-GaMD simulations and SEC indicate that free dimeric forms explore highly heterogeneous, non-productive conformations in solution. Stable, functional dimerization strictly requires the DNA marbox to act as an organizing scaffold, supporting a monomer-driven binding pathway. In this framework, the C-terminal extension does not primarily drive initial DNA affinity, but rather ensures the correct quaternary organization, inter-monomer stability, and precise DNA bending angle upon binding.

Ultimately, the ability of a single HTH motif to cooperatively engage two half-sites offers a compelling structural rationale for the existence of naturally occurring short AraC/XylS-like proteins that lack the full two-motif architecture. This capacity of a single HTH motif to dimerize and regulate transcription provides strong biochemical support for a gene-duplication origin of modern, multidomain AraC/XylS transcription factors, a hypothesis further supported by the existence of predicted proteins harbouring a single MarA-like HTH motif in extant genomes.

These insights redefine the minimal structural requirements for marbox recognition and also establish a foundation for engineering compact, highly specific transcriptional regulators for synthetic biology and antimicrobial applications.

## Materials and Methods

### Cloning and site-directed mutagenesis

Wild-type MarA (residues 1–127, N-terminal hexahistidine tagged) was cloned into pET15b (*pmarA) as* explained in (Corbella et al. 2025). Both truncations, MarA64 (residues 1-64) and MarA57 (residues 1-57), were obtained by site-directed mutagenesis carried out using the Quick-Change Lightning SDM Kit (Agilent), the plasmid p*marA* as template and primers 5’CGTAAGATGACGTAAATCGCGCAAAAGCTG3’ (forward) and 5’GCTTTTGCGCGATTTACGTCATCTTACGGCTG3’ (reverse) for obtaining MarA64 and 5’CATTAGGCCAATACTAACGCAGCCGTAAGATG3’ (forward) and 5’CTTACGGCTGCGTTAGTATTGGCCTAATGAATG3’ (reverse) for obtaining MarA57. Sanger sequencing confirmed the stop codon in all constructs.

### Protein expression and purification

WT MarA, MarA64 and MarA57, were expressed in T7 express *E. coli* cells (New England Biolabs) grown to OD600≈0.5 and induced with 0.5 mM IPTG for 3 h at 37°C. The cells were harvested by centrifugation (5000g, 15 min, at 4°C), resuspended in 50 mM Tris–HCl pH 8, 1M NaCl, 0.03 mg/mL DNase I (Sigma), supplemented with a protease inhibitor cocktail pill (Roche), and lysed by sonication. Inclusion bodies containing MarA were collected by centrifugation at 75,000g for 30 min and washed with 50 mM Tris–HCl pH 8, 1M NaCl, 2M urea and centrifuged at 75,000g for 30 min. Inclusion bodies were solubilized with 50 mM Tris–HCl pH 8, 1M NaCl, 7M urea and isolated by high-speed centrifugation (75,000g, 30 min). The supernatant was loaded in a 0.1-ml His-SpinTrap centrifugal column (Cytiva), equilibrated with buffer A (50 mM Tris–HCl pH 8, 1M NaCl, 7M urea, 50 mM imidazole) and eluted with 0.3 M imidazole-containing buffer A. Purified proteins were buffer exchanged into 50 mM Hepes pH 8, 1M NaCl by using PD SpinTrap G-25 centrifugal columns (Cytiva).

### Electrophoretic mobility shift assays

For electrophoretic mobility shift assay (EMSA) experiments, the 200-bp DNA fragments containing the marRAB or acrAB marboxes were prepared by PCR amplification using the primers listed in Table S1 and *E. coli* MG1655 genomic DNA as template. The 30-bp complementary DNA fragments corresponding to the marRAB or acrAB marboxes with five nucleotides in each flank were purchased from Eurogentec (Seraing, Belgium; Table S1). The DNA fragments (15 and 60 nM when using 200-bp and 30-bp DNA fragments, respectively) were incubated with MarA in a buffer containing 20 mM Hepes pH 8, 10 mM MgCl_2_, 100 mM EDTA, and 0.3 mg/mL BSA. 1 - 2.5 μM of purified *E. coli* MarA WT or 5 – 6 μM of the truncations were added and the mix was incubated for 20 min at 37°C (final volume 10 μL). The samples were mixed with 6X EMSA gel-loading solution (component D, Electrophoretic Mobility Shift Assay (EMSA) Kit, Invitrogen) and loaded onto an 8% native acrylamide gel and visualized with SYBR green (Invitrogen).

### Size exclusion chromatography

Size-exclusion chromatography was performed using a Nexera ultra-high-performance liquid chromatography (UHPLC) system (Shimadzu, Kyoto, Japan) equipped with an SPD-40 UV-Vis detector and a bioZen SEC-2 column (150 × 4.6 mm, 1.8 μm; Phenomenex, Torrance, CA, USA). The samples were eluted under isocratic conditions using 0.1 M ammonium acetate, pH 6.8 for 30 minutes at a flow rate of 0.2 mL/min. 5 µ L of 0.1 mg/ml of WT MarA, MarA64, MarA57, and a buffer blank were monitored by UV absorbance at 220 nm. Buffer was subtracted and data were normalised to 1 for each run.

### System preparation for MD simulations

Starting coordinates for all the simulations in this work were generated using AlphaFold3 (Abramson et al. 2024) (ipTM and pTM scores are reported for each model). Two sets of simulations were performed, one set describing free MarA truncations (MarA57 and MarA64), and a second describing dimer–DNA complexes (MarA57-*mar* and MarA64-*mar*), using the 30-bp *mar* fragment 5’GAACCGAT**TTA**GCAAAAC**GTGG**CATCGGTC3’). Relevant data concerning the molecular dynamics simulations are available for download from Zenodo, https://doi.org/10.5281/zenodo.22880098.

### Classical MD simulations

Conventional molecular dynamics (cMD) simulations were performed following the same protocol as our prior studies of MarA-DNA complexes (Corbella et al. 2021; Corbella et al. 2025). In brief, all simulations were performed using the Amber ff14SB force field (Maier et al. 2015) to describe MarA64 and MarA57, the Parmbsc1 force field to describe the DNA (Ivani et al. 2016), and the CUDA version of the PMEMD module (Götz et al. 2012) of the AMBER 20 simulation package (D.A. Case et al. 2020). Three independent production runs of 2.5 μs of length each with different initial velocities were performed for each system. Further simulation details can be found in (Corbella et al. 2021).

### PPI-GaMD simulations

Gaussian accelerated MD (GaMD) (Miao et al. 2015) enhances conformational sampling by applying a non-negative, Gaussian-distributed boost potential to smooth the system’s potential energy surface (PES). This effectively lowers energy barriers and accelerates transitions between distinct low-energy minima. Extending this concept, the PPI-GaMD approach specifically investigates protein-protein interactions (PPIs) by selectively boosting the intermolecular interaction energies (both electrostatic and van der Waals) to promote protein dissociation. Concurrently, a secondary boost is applied to the remaining potential energy of the system to model the system’s flexibility and encourage subsequent rebinding events (Wang and Miao 2022).

In this work PPI-GaMD was applied to simulations of both MarA64 and MarA57 free dimeric systems, and the same AMBER ff14SB force field (Maier et al. 2015) was used here for the protein. The system was neutralized by adding counter ions and immersing in a cubic TIP4P2015 water box (Jorgensen et al. 1983), which was extended for 18 Å from the protein−protein complex surface. This water model was chosen here because it was shown to be more accurate in calculating kinetic parameters (Pan et al. 2019; Wang and Miao 2022).

The simulation protocol used was extracted from the original PPI-GaMD article (Wang and Miao 2022). The systems were first energy minimized with 1.0 kcal/mol/Å^2^ constraints on the heavy atoms of the proteins, including the steepest descent minimization for 50,000 steps and conjugate gradient minimization for 50,000 steps. The system was then heated from 0 to 300 K for 200 ps. It was further equilibrated using the NVT ensemble at 310 K for 200 ps and the NPT ensemble at 300 K and 1 bar for 1 ns with 1 kcal/mol/Å^2^ constraints on the heavy atoms of the protein, followed by 2 ns short cMD without any constraint. The PPI-GaMD simulations proceeded with 14 ns short cMD to collect the potential statistics, 46 ns PPI-GaMD equilibration, during which time the boost potential was updated every 5.0 ns, and finally six independent 2 ms production runs. The threshold energies for applying the boost potentials were all set to the upper bound (i.e., *E* = *V_min_* + 1/*k*) (Miao et al. 2020; Wang and Miao 2020). In order to observe protein dissociation during PPI-GaMD equilibration while keeping the boost potential as low as possible for accurate energetic reweighting, the σ_0P_ and σ_0D_ parameters were set to 2.9 and 7.0 kcal/mol, respectively, in the final PPI-GaMD simulations.

### MD simulations analysis

All analysis of MD simulations were performed using CPPTRAJ (Roe and Cheatham 2013). The most-populated structures were calculated by performing agglomerative hierarchical clustering on the root mean square deviation (RMSD) of Cα-atoms to yield ten discrete clusters. The centroid of the most-populated cluster was then selected for further

characterization. Secondary structural propensities for all residues of free MarA were calculated using the DSSP method of Kabsch and Sander (Kabsch and Sander 1983). T-shaped p-p interaction parameters, R_c_ and g, were calculated as the distance between the COM of all the heavy atoms of each ring and the angle between the normal vectors of each ring plane. The DNA bending angle was calculated as the angle between two vectors along both halves of the double helix sequence (G8/C53 to C4/G57 and C34/27 to A22/T39). An interaction was defined as a hydrogen bond if the distance between the donor and the acceptor on the protein/DNA fell within 3.5 Å.

### Reporter gene assay

*E. coli* T7 Express cells carrying a pACYC177 plasmid composed of the *E. coli acrAB* promoter followed by the GFP protein (named pACYC177-RS) was co-transformed with the pET-15b empty vector, or harbouring the WT MarA, or MarA64, or MarA57. Cultures were grown up to OD600 = 0.5 and induced with 0.5 mM IPTG for 30 min at 37°C. The cells were harvested by centrifugation, the pellet was resuspended with MOPS minimal medium, and 190 μL of all the strains were placed in a 96-well black/clear bottom plate (Thermo Fisher) and the OD600 and fluorescence was measured. The fluorescence signals were normalized using the number of cells for every culture. Data presented are the mean of at least three biological replicates. Statistical comparisons between T7 Express cells, WT MarA, MarA64, MarA57 and the pET15b empty-vector control were performed using an unpaired, two-tailed Student’s t-test in GraphPad Prism. A P value < 0.05 was considered statistically significant.

### Growth curve under antibiotic stress

*E. coli* T7 Express cells were transformed with pET15b empty vector, or pET15b encoding WT MarA (p*marA*), MarA64, or MarA57. Cultures were grown to OD600 = 0.5 and induced with 0.5 mM IPTG for 3 h at 37°C prior to the growth-curve assay, to allow adaptation. 0.5 mM IPTG and 50 µg/mL ampicillin were maintained throughout the growth-curve assay to sustain MarA overexpression and plasmid selection, respectively. Following induction, cultures were exposed to erythromycin at two concentrations (4 and 256 µg/mL) or to gentamicin (2, 4 and 256 µg/mL) as a non-AcrAB-TolC-substrate control. Growth was monitored by absorbance at 600 nm using a Tecan Infinite 200 Pro microplate reader in Greiner 96-well flat-bottom transparent polystyrene plates. Plates were incubated at 37°C with absorbance reads taken every 10 min over a total duration of 16 h. Each cycle included 10 s of orbital shaking (2 mm amplitude) immediately before the read. Data presented are the mean of at least two biological replicates. Statistical comparisons between pET15b empty vector, MarA64, MarA57 and WT MarA were performed using an unpaired, two-tailed Student’s t-test in GraphPad Prism. A P value < 0.05 was considered statistically significant.

### Computational search for single MarA-like HTH proteins

MarA, Rob and SoxS sequences from *E. coli* K-12 MG1655 (NP_416048.2, NP_418813.1 and NP_418486.1) were used as queries for a BLASTP search excluding *Enterobacteriaceae* (taxID: 543) with a 1e-40 limiting e-value (Altschul et al. 1997). For each query, the sequences for the top 7 hits with query coverage above 90% were downloaded. The *E. coli* MarA, Rob and SoxS sequences and those of identified homologs were aligned using the M-COFFEE server with default parameters, and the multiple sequence alignment was manually trimmed with BioEdit to include only the region spanning the MarA HTH1 motif (Hall 1999; Moretti et al. 2007). The resulting trimmed alignment was used to create a HMM profile with the hmmbuild program of the HMMER suite (Eddy 2011), and the profile was used to search the UniProt database (UniProt (2025_01)) with hmmsearch, filtering for Bacteria (taxID: 2) and with otherwise default parameters (Rajkovic et al. 2026).

### Artificial intelligence statement

During the preparation of this work, the author(s) used Claude (*Anthropic)* to assist with literature search and to improve the English language and style of the manuscript. After using this tool, the author(s) reviewed and edited the content as needed and take full responsibility for the content of the publication.

### Author contributions

**Marina Corbella:** Conceptualization; methodology; investigation; visualization; supervision; writing – original draft; writing – review and editing. **Ariadna Serrano:** investigation. **Roberto Bello-Madruga**: Investigation; visualization; writing – review and editing. **Marc Torrent Burgas:** investigation; writing – review and editing. **Ivan Erill:** methodology; investigation; visualization; writing – original draft; writing – review and editing. **Jessica M. A. Blair:** Conceptualization; methodology; supervision; writing – original draft; writing – review and editing. **Enea Sancho-Vaello:** Conceptualization; methodology; investigation; visualization; supervision; writing – original draft; writing – review and editing.

## Supporting information

Supplemental figures S1-S9, supplemental tables S1-S3, supplementary references

## Acknowledgements

We gratefully acknowledge Professor Carles Curutchet for valuable scientific discussions. We thank Dr. Vito Ricci and Professor Laura Piddock for sharing the pMW82 plasmid harbouring the *E. coli acrAB*-GFP reporter system. This work was supported by the European Union’s Horizon 2020 Research and Innovation Programme through the Marie Sklodowska-Curie Grant, grant number 839036, the Beatriz Galindo Programme (grant agreement BG22/00118) from the Spanish Ministry of Science and Innovation, given to E.S.-V., and the Universitat Autònoma de Barcelona (grant agreement PD619842/D040601). Marina Corbella gratefully acknowledges RES resources provided by BSC in MareNostrum5 to BCV-2025-3-0015. Ivan Erill acknowledges the Spanish Ministry of Science, Innovation and Universities (MICIU) under grants PID2023-152240OB-C21 and PID2024-156292OB-I00.

