## Supplemental figures S1-S9, supplemental tables S1-S3, supplementary references for "Seven amino acids gate transcriptional activation by a minimal MarA Helix-turn-helix DNA-binding domain"

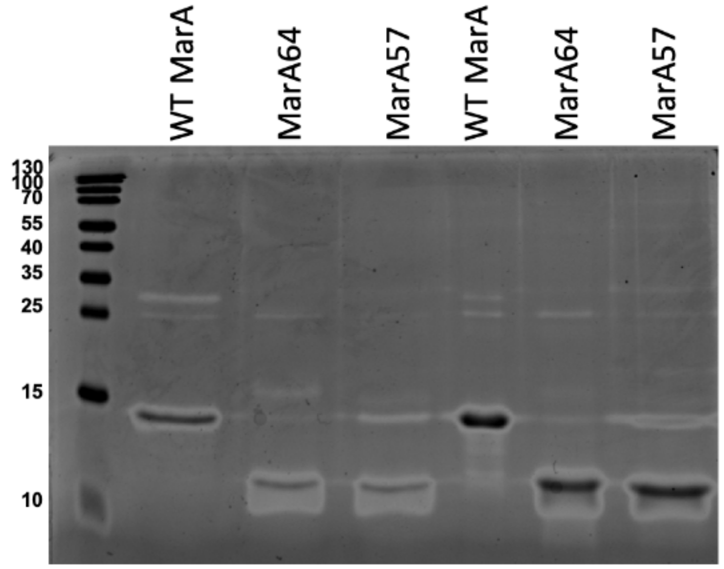

**Supplemental Figure S1.** SDS-PAGE of two independent purifications of WT MarA, MarA64, and MarA57. A band consistent with an SDS-resistant MarA57 dimer is observed at the same position as the WT MarA monomer band. The three constructs (WT MarA, MW 17.35 kDa, pI 9.52; MarA64, MW 9,757 Da, pI 9.86; MarA57, MW 8,884 Da, pI 8.25) are all cationic, which is known to cause anomalously fast migration on SDS-PAGE. Construct identity was confirmed by DNA sequencing, and the absence of WT MarA contamination in the truncated constructs was confirmed by HPLC-MS.

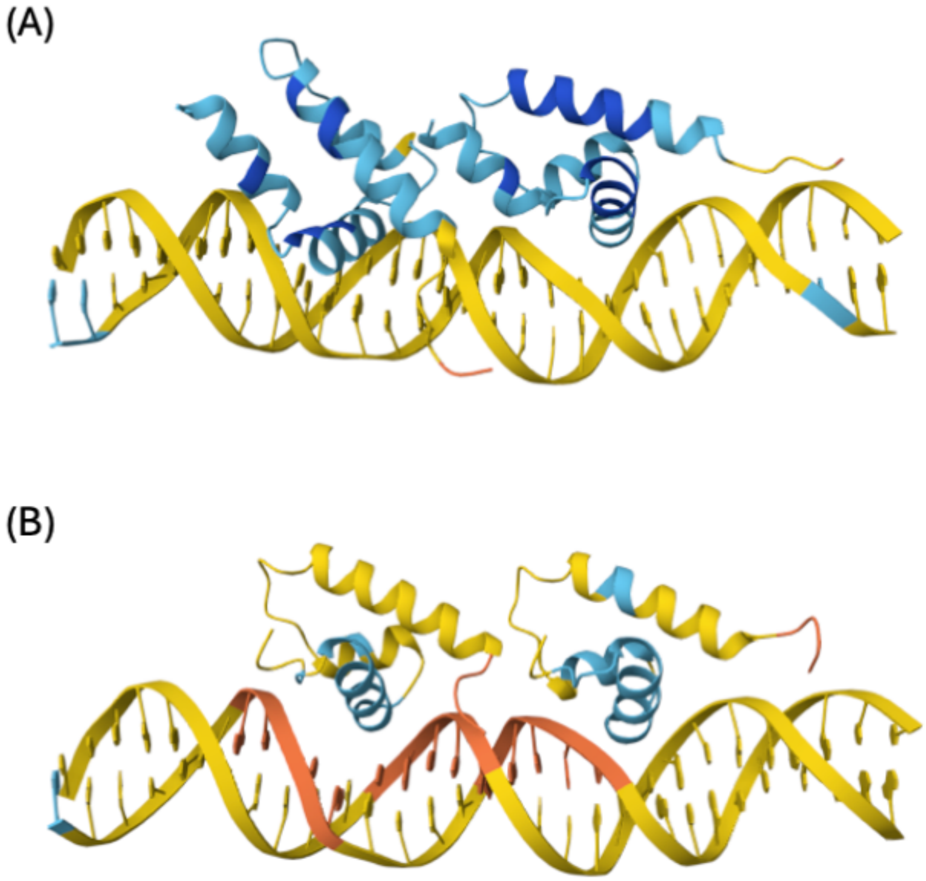

**Supplemental Figure S2.** AlphaFold structure prediction of **(A)** MarA64 dimer-DNA and **(B)** MarA57 dimer-DNA complexes. Predictions are coloured from red (worst) to dark blue (best) according to pLDDT values. Overall structure prediction is far beyond the confidence threshold, ipTM = 0.34, pTM = 0.48 and ipTM = 0.28, pTM = 0.36, respectively.[1]

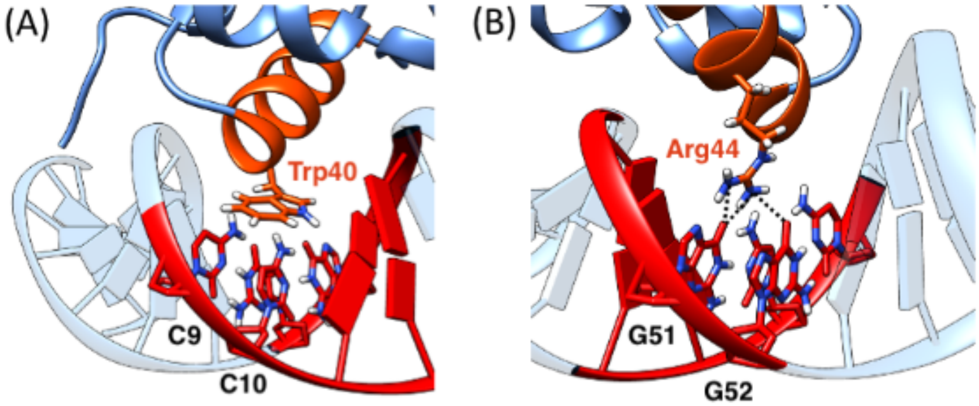

**Supplemental Figure S3.** Key interactions established between Trp40 and Arg44 from helix 3 of monomer A of MarA64 and the corresponding nucleobases within A-box. **(A)** T-shaped $\pi$–$\pi$ interactions with C9 and C10 (Table S2), and **(B)** stable hydrogen bonds with G51 and G52 (Table S3). Structures were selected based on visual examination of the trajectories.

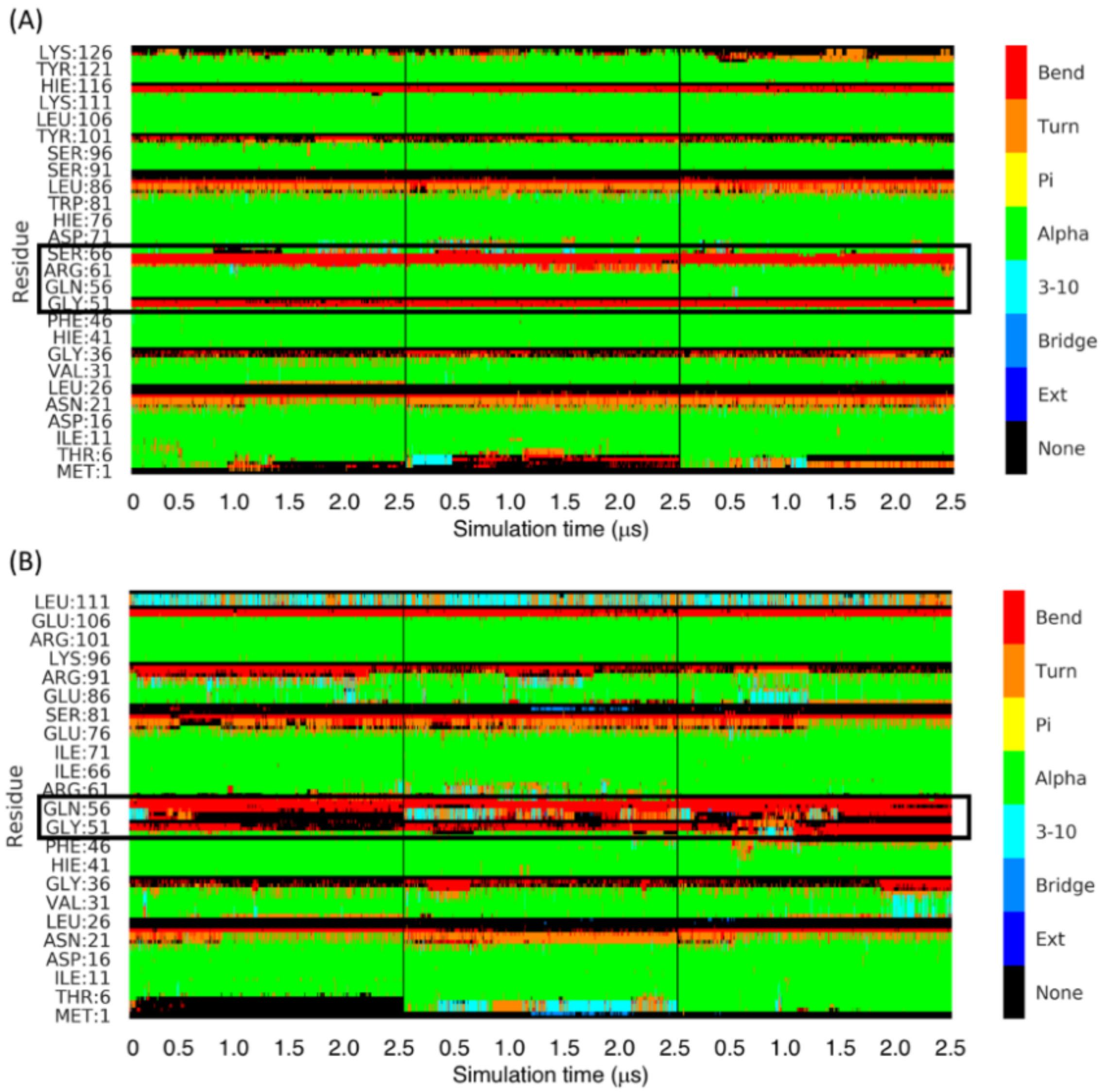

**Supplemental Figure S4.** Time evolution of secondary structure composition of **(A)** MarA64 and **(B)** MarA57 dimers in complex with DNA. It is highlighted with a black box, the region corresponding to the C-terminal IRSRKMT sequence truncation as well as a few residues before. Data was extracted for 3 independent 2.5 μs MD simulations.

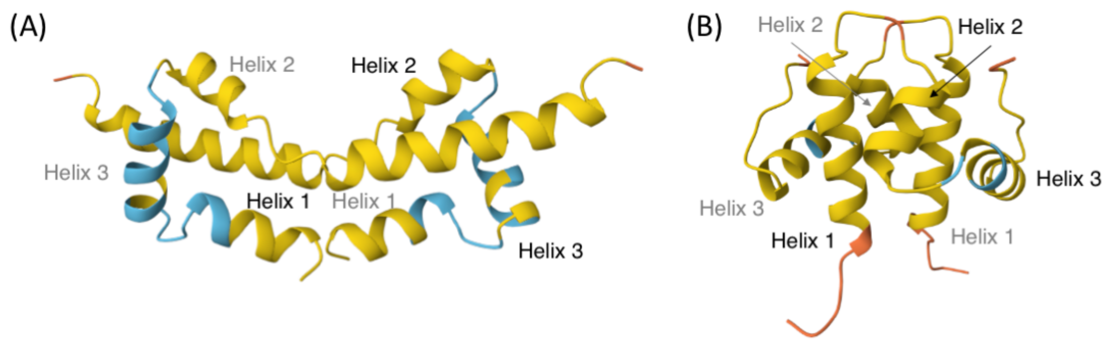

**Supplemental Figure S5.** AlphaFold structure prediction of **(A)** MarA64 and **(B)** MarA57 free dimers. Predictions are coloured from red (worst) to dark blue (best) according to pLDDT values. Overall structure prediction is far beyond the confidence threshold, ipTM = 0.25, pTM = 0.35 and ipTM = 0.15, pTM = 0.28, respectively.[1]

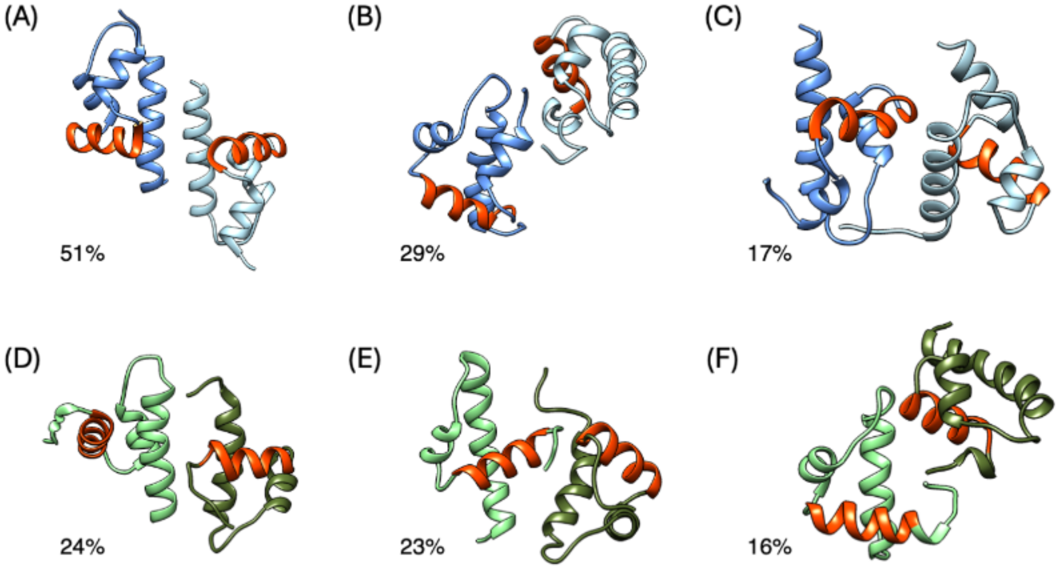

**Supplemental Figure S6.** Structures of the most populated clusters of MarA64 (top row) and MarA57 (bottom row). Each monomer is colored in a different shade of blue (MarA64) or green (MarA57), while helixes 3 are colored in orange. Data was extracted along 6 independent 2.5 μs PPI-GaMD simulations.

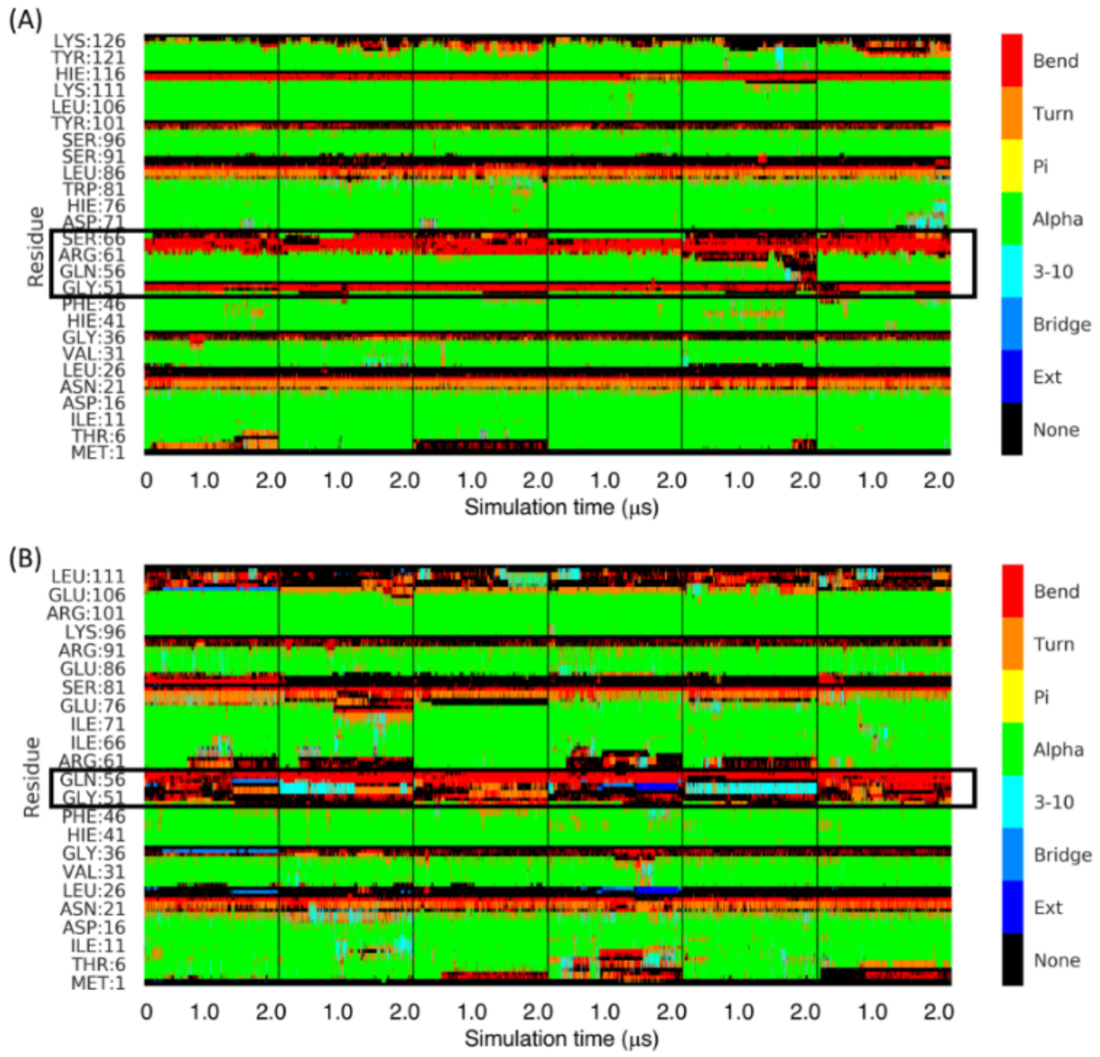

**Supplemental Figure S7.** Time evolution of secondary structure composition of **(A)** MarA64 and **(B)** MarA57 free dimers. It is highlighted with a black box, the region corresponding to the C-terminal IRSRKMT sequence truncation as well as a few residues before. Data was extracted for 6 independent 2.0 μs MD simulations.

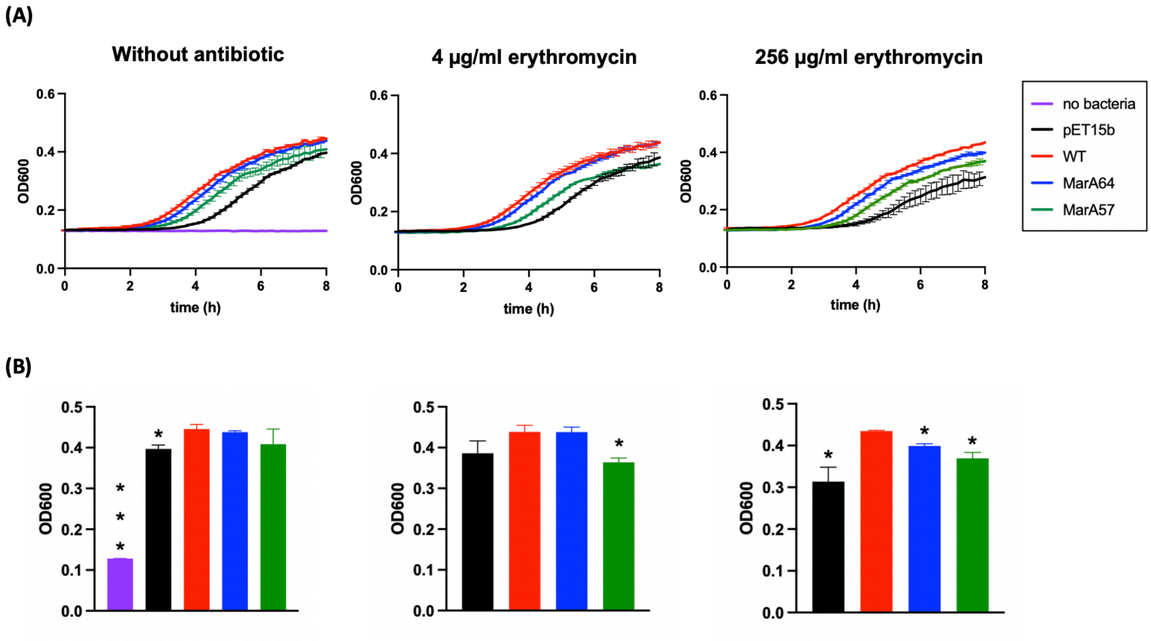

**Supplemental Figure S8.** Growth of MarA constructs under erythromycin stress. **(A)** *E. coli* T7 Express cultures carrying pET15b empty vector (black), WT MarA (red), MarA64 (blue), or MarA57 (green) were grown in the absence, low (4 µg/mL) and high (256 µg/mL) concentration of erythromycin. The experiment was performed in duplicate, and the standard error of the mean (SEM) is shown for each curve. **(B)** Bars show strain growth at a fixed timepoint (8 h). The asterisks indicate statistically significant differences between WT MarA and the rest of the strains (tested by a Student's t test).

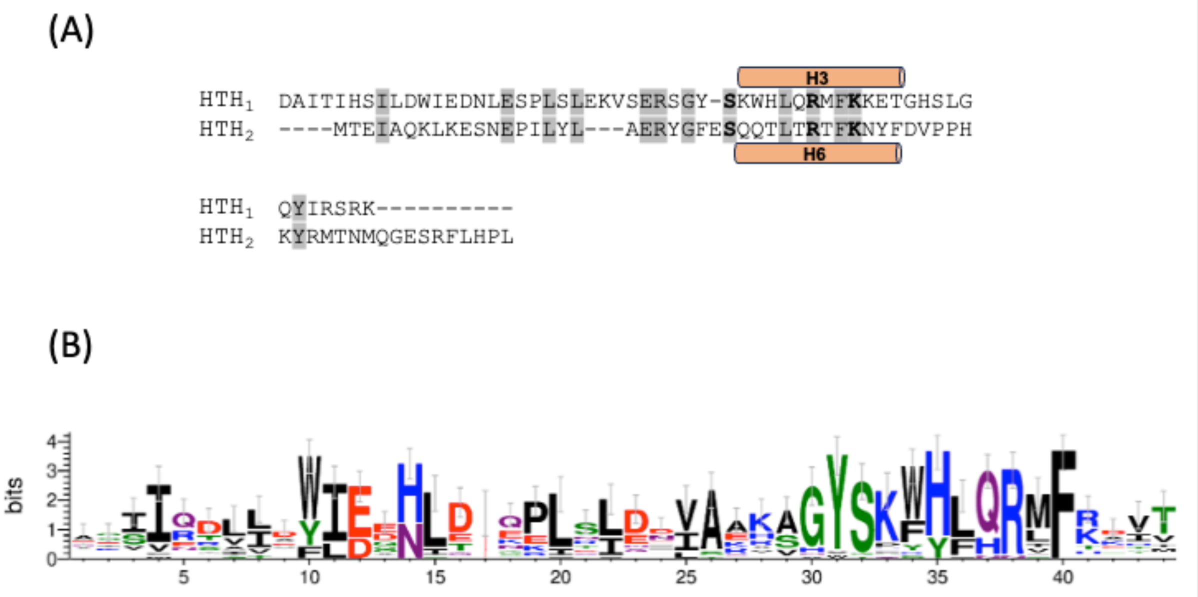

**Supplemental Figure S9: (A)** Sequence alignment of the N- and C-terminal HTH domains of MarA (HTH1 and HTH2), with DNA-contacting helices (H3 and H6) indicated by orange bars alongside the corresponding sequence. Conserved residues are shaded grey, and residues critical for DNA binding are shown in bold. **(B)** Sequence logo for the HMM profile of the MarA HTH1, spanning MarA helices H1, H2 and H3.

**Supplemental Table S1.** Sequences of the primer pairs used to generate the ~200-bp DNA fragments containing the marbox, as well as the 30-bp oligonucleotides used directly as probes in EMSAs.

| Primer name | Primer sequence (5´- 3´) |
|---|---|
| **Amplification of the ~200-bp DNA fragments used in the EMSAs** | |
| *acrAB*_long_prom.FOR | GCTTTTGCAATCTCGCCCAGC |
| *acrAB*_long_prom.REV | GTCCGATTTCAAATTGGTCAATGG |
| *marRAB*_long_prom.FOR | GTTATCCTGTGTATCTGGGTTATCAGCG |
| *marRAB*_long_prom.REV | GTTGCCCTGGCAAGTAATTAGTTGC |
| **30-bp DNA fragments used in the EMSAs** | |
| *acrAB*_short_prom.FOR | CTTCTTGTTTGGTTTTTCGTGCCATATGTTC |
| *acrAB*_short_prom.REV | GAACATATGGCACGAAAAACCAAACAAGAAG |
| *marRAB*_short_prom.FOR | GAACCGATTTAGCAAAACGTGGCATCGGTC |
| *marRAB*_short_prom.REV | GACCGATGCCACGTTTTGCTAAATCGGTTC |
| non_specific.FOR | CGATTGGCTACATCCAACAACCTGGAATCG |
| non_specific.REV | CGATTCCAGGTTGTTGGATGTAGCCAATCG |

**Supplemental Table S2.** Average distances between the Trp41 and C9/C10 and C19/G18 ring centroids ($R_c$), and angles ($\gamma$) between normal vectors of each ring plane, defining T-shaped $\pi$–$\pi$ interactions between these residues.[a]

| System | Parameters | A Box | | B Box | |
|---|---|---|---|---|---|
| | | **C9** | **C10** | **C19** | **G18** |
| **MarA64** | $R_c$ | 5.3 ± 0.6 | 5.3 ± 0.4 | 6.5 ± 2.0 | 7.0 ± 1.6 |
| | $\gamma$ | 110.2 ± 11.3 | 112.1 ± 12.2 | 98.2 ± 34.0 | 99.4 ± 32.5 |
| **MarA57** | $R_c$ | 6.0 ± 1.8 | 6.2 ± 1.7 | 7.4 ± 2.4 | 8.6 ± 2.1 |
| | $\gamma$ | 73.7 ± 19.7 | 75.2 ± 17.6 | 93.7 ± 29.6 | 91.1 ± 29.5 |

[a] A T-shaped $\pi$–$\pi$ interactions between two rings is defined as $R_c$ between 4.5 and 7.5 Å, and $\gamma$ between 50 and 90°.[2–4]

**Supplemental Table S3.** Stability of A- and B-box hydrogen-bonding interactions between Arg44 of each monomer of MarA64 and MarA57 along 3 independent 2.5 μs MD simulations. Values shown as % of simulation time.

| Arg44 | DNA | A Box | | DNA | B box | |
|---|---|---|---|---|---|---|
| | | **MarA64** | **MarA57** | | **MarA64** | **MarA57** |
| NH1 | G51 O6 | 15.2 | 3.3 | G18 O6 | 67.5 | 16.3 |
| NH2 | G51 O6 | 35.4 | 18.7 | G18 O6 | 26.9 | 24.1 |
| NH1 | G52 O6 | 35.4 | 16.1 | G42 O6 | 1.4 | 39.9 |
| NH2 | G52 O6 | 31.1 | 43.0 | G42 O6 | 42.9 | 44.4 |
| NH1 | G8 O6 | 20.9 | 53.5 | | | |
